# Propofol Anesthesia Concentration Rather Than Abrupt Behavioral Unresponsiveness Linearly Degrades Responses in the Rat Primary Auditory Cortex

**DOI:** 10.1101/2020.10.28.359323

**Authors:** Lottem Bergman, Aaron J Krom, Yaniv Sela, Amit Marmelshtein, Hanna Hayat, Noa Regev, Yuval Nir

## Abstract

Despite extensive knowledge of its molecular and cellular effects, how anesthesia affects sensory processing remains poorly understood. In particular, it remains unclear whether anesthesia modestly or robustly degrades activity in primary sensory regions, and whether such changes are linked to anesthesia drug concentration vs. behavioral unresponsiveness, since these are typically confounded. Here, we employed slow gradual intravenous propofol anesthesia induction together with auditory stimulation and intermittent assessment of behavioral responsiveness while recording epidural EEG, and neuronal spiking activity in the primary auditory cortex (PAC) of eight rats. We found that all main components of neuronal activity including spontaneous firing rates, onset response magnitudes, onset response latencies, post-onset neuronal silence duration, and late-locking to 40Hz click-trains, gradually deteriorated in a dose-dependent manner with increasing anesthesia levels without showing abrupt changes around loss of righting reflex or other time-points. Thus, the dominant factor affecting PAC responses is the anesthesia drug concentration rather than any sudden, dichotomous behavioral state changes. Our findings explain a wide array of seemingly conflicting results in the literature that, depending on the precise definition of wakefulness (vigilant vs. drowsy) and anesthesia (light vs. deep/surgical), report a spectrum of effects in primary regions ranging from minimal to dramatic differences.

## Introduction

Anesthesia constitutes a powerful model for inducing changes in consciousness and sensory perception in a controlled, reversible manner with fine temporal resolution (Alkire et al. 2008; Mashour et al. 2020). Despite the widespread use of anesthesia since the 19^th^ century (Morton 1847), and considerable progress in understanding its molecular and cellular effects (Rudolph and Antkowiak 2004; Franks 2008; Brown et al. 2011), how anesthesia affects sensory responses remains unclear. Specifically, a controversy persists about the extent to which anesthesia affects activity in primary sensory cortices.

Most knowledge on responses in primary sensory regions, and the principles that organize their activities (e.g. tuning-curves, maps) were obtained in anesthetized animals (Mountcastle 1957; Hubel and Wiesel 1959; Merzenich et al. 1975). Studies directly comparing sensory responses in primary visual and somatosensory cortices across wakefulness and anesthesia revealed degraded responses during anesthesia (Wörgötter et al. 1998; Imas et al. 2005; Ferezou et al. 2006; Constantinople and Bruno 2011; Haider et al. 2013). However, the absence of active sensing (e.g., eye movements, whisking) in anesthesia, which is known to play a key role in these modalities (Kleinfeld et al. 2006), could potentially contribute to such differences and therefore limits the ability to attribute changes in sensory responses to anesthesia drug concentration or behavioral states changes. The controversy about whether anesthesia robustly degrades responses in primary sensory regions also exists in the auditory domain, even though auditory processing is largely passive. While some studies reported robust deterioration in response magnitude in the primary auditory cortex (PAC) (Katsuki, Murat, et al. 1959; Katsuki, Watanaba, et al. 1959; Capsius and Leppelsack 1996; Gaese and Ostwald 2001; Banks et al. 2018; Wang 2018; Du et al. 2020), other studies only report modest changes in response magnitude, if any (Davis et al. 2007; Raz et al. 2014; Krom et al. 2020).

Most previous studies that compared sensory processing across wakefulness and anesthesia employed deep anesthesia at a narrow range of deep surgical levels (Schwender et al. 1993; Capsius and Leppelsack 1996; Plourde 1996; Gaese and Ostwald 2001; Heinke et al. 2004; Ferezou et al. 2006; Sellers et al. 2013; Wang et al. 2018), and most animal studies did not behaviorally identify the precise moment of behavioral states changes (e.g. loss of righting reflex, or EEG signatures of loss of consciousness such as appearance of alpha-band enhancement (Purdon et al. 2013)). The main limitation in nearly all previous studies is that anesthesia drug concentration and behavioral (un)responsiveness were confounded and compared in a dichotomic manner in two experimental conditions (drug-free wakefulness vs. anesthesia with behavioral unresponsiveness). It remains unknown if the reported changes in sensory responses stem from the amount of anesthetic given, or alternatively from behavioral state changes.

Our goal in this study was to go beyond these limitations by using slow gradual anesthesia induction together with intermittent behavioral assessment and continuous epidural EEG recording, while following the precise dynamics of auditory responses at the neuronal level. We sought to examine whether anesthesia modestly or robustly degrades activity in PAC, and whether observed changes are linked to depth of anesthesia (i.e. amount of anesthetic given) vs. behavioral unresponsiveness. We used auditory 40Hz click-trains as stimuli, which are known to be highly effective in revealing changes between wakefulness and anesthesia across species. PAC responses to 40Hz click-trains yield excellent signal-to-noise ratio (SNR) that allow comparing sensory responses in short time intervals (Plourde 1996; Conti et al. 1999; de Bruin et al. 1999; Santarelli et al. 2003; Anderson et al. 2006; Plourde et al. 2008; Vohs et al. 2012; Ma et al. 2013; Sivarao 2015; Banks et al. 2018; Krom et al. 2020). We find that all main components of neuronal activity in PAC gradually deteriorate in a dose-dependent manner with increasing anesthesia levels without showing abrupt changes around loss of righting reflex or other time-points. This indicates that the dominant factor affecting PAC responses is the anesthesia drug concentration rather than behavioral state changes.

## Materials and Methods

### Animals

Experiments were performed in eight adult male Long-Evans (250-350g) rats individually housed in a transparent Perspex cage, with food and water available *ad libitum*. Ambient temperature was kept at 22-23° Celsius, with a light-dark cycle of 12:12h starting at 10AM. All experimental procedures including animal handling, surgery and experiments were approved and supervised by the Institutional Animal Care and Use Committee (IACUC) of Tel Aviv University (approval M-15-066).

### Surgery and electrode implantation

During surgery, animals were anesthetized using Isoflurane (4-5% induction, 1-2% maintenance) and placed in a stereotactic frame (David Kopf Instruments; CA, USA), while maintaining constant body temperature using a heating pad (Harvard Apparatus, MA, USA). Chronic central venous catheter (Stewart 1981),(Braintree R-JVC-M37-R 120mm) was implanted in the jugular vein, and tunneled subcutaneously to the dorsal neck, and closed with Heparin-Lock (Luo et al. 2000). The head was shaved and liquid Viscotear gel was applied to protect the eyes. The skin was incised to expose the skull. Frontal and parietal screws (1mm in diameter) were placed on the left side of the skull for electroencephalogram (EEG) recording, two more screws above the cerebellum were used as reference and ground, and an anchor screw located above the right frontal lobe for stabilization (Figure 1A). Two stainless-steel wires were bilaterally inserted to the neck muscles to measure electromyography (EMG). The EEG and the EMG wires were soldered into a headstage connector (Omnetics Connector Corporation; MN, USA).

**Figure 1.**
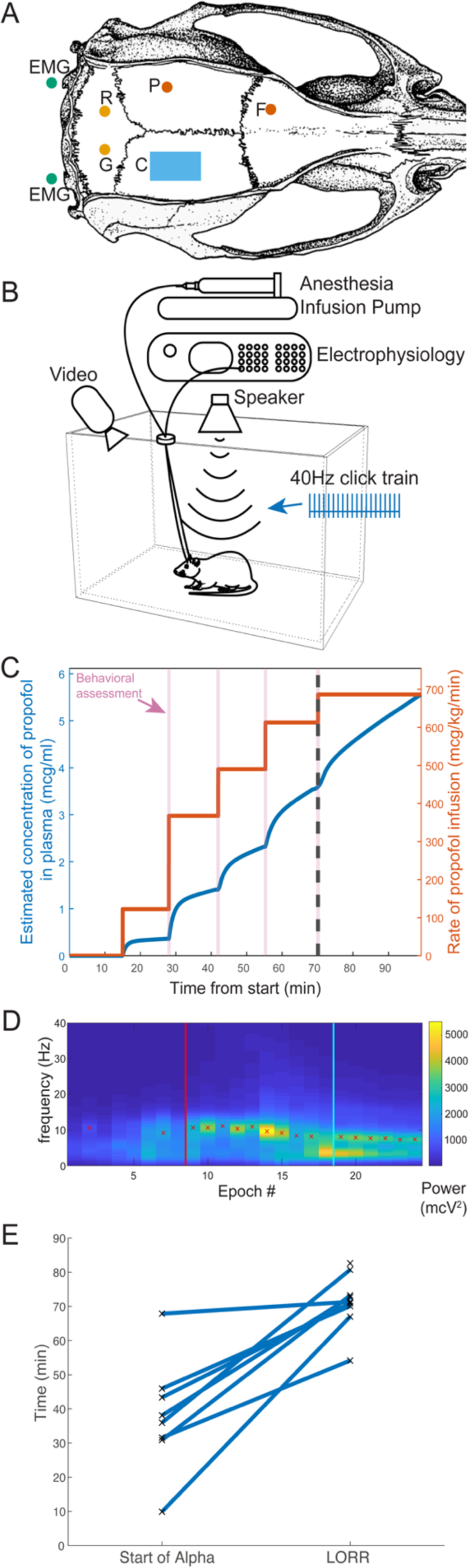
Experimental design. (A) Surgical plan includes frontal (“F”) and parietal (“P”) EEG screws (red circles), ground (“G”) and reference (“R”) screws (orange circles) above cerebellum, and two wires (green circles) for recording EMG from neck muscles. Microwire arrays are implanted diagonally through a dorsal craniotomy (C, blue rectangle) to target the primary auditory cortex. (B) Experimental setup: rats were placed individually inside a sound-attenuating chamber, while auditory stimuli were delivered free-field from a speaker above. Electrophysiology data were continuously collected while rats were gradually anesthetized with propofol via a computer-controlled syringe pump. (C) Anesthesia dynamics over time (xaxis) in a representative experiment: propofol rate infused by pump (orange, right y-axis), estimated propofol concentration in plasma (blue, left y-axis), and intermittent behavioral assessments every 12 min (pink vertical lines; dashed black line marks loss of righting reflex, LORR). (D) Representative time-frequency EEG dynamics (spectrogram) during an experimental session, demonstrating appearance of alpha activity (warm colors) starting around Epoch 9, and LORR after Epoch 18. Red crosses indicate identified EEG alpha peaks, and red vertical line indicates start of alpha enhancement (Methods) Cyan vertical line marks LORR. (E) time to start of frontal EEG alpha enhancement vs. LORR for all experimental sessions, demonstrating that alpha enhancement (when detectable) precedes LORR.

A small craniotomy was performed and dura was carefully reflected under microscopic control. Rats were implanted with a 32-channel microwire array (Tucker-Davis Technologies Inc, ‘TDT’, Alachua, FL, USA), with wire diameter of 50 µm, and tip angle of 45°. The 32 wires were ordered in 4 rows of eight channels, with 375 µm separation along the medial-lateral axis between rows, and 250 µm separation along the anterior-posterior axis within each row. Two rows of 8 long microwires each (length: 9.6 mm and 9.4 mm) targeted perirhinal cortex (not included in this study), while two rows of 8 short microwires each (length: 7.7 mm and 7.2 mm) targeted primary auditory cortex. Implantation was performed diagonally (insertion point P: 3.6 mm, L: 4.2 mm relative to Bregma and inserted to a depth of 8mm) with 22° angle (Figure 1A). DiI fluorescent dye (DiIC18 (3), Invitrogen) was applied on the microwires to facilitate subsequent histological identification. After implantation, the craniotomy was covered with silicone (Kwik-Sil; World Precision Instruments, FL, USA) followed by Fusio (Pentron, Czech Republic). Dental cement was then used to cover all screws and wires.

### Antibiotic prophylaxis and peri/postoperative analgesia

Antibiotic prophylaxis (cefazolin, 20 mg/kg i.m.) was administered before incision, and local chloramphenicol 3% ointment applied at the end of surgery. Carprofen (5 mg/kg) and dexamethasone (0.5 mg/kg) were administrated intraperitoneally after anesthesia induction, and the surgical site was anesthetized subcutaneously with lidocaine (3mg/kg) before incision. Postoperative analgesia was provided by injecting buprenorphine systemically (0.025 mg/kg s.c.) as the rat started showing signs of awakening from anesthesia. In the days following surgery, dexamethasone (1.3 mg/kg) was given with food to reduce pain, edema and scare tissue around implanted electrodes.

### Electrophysiology

All electrophysiological data were acquired continuously using an RZ2 processor (TDT). We recorded extracellular spiking activity (24.4 kHz sampling rate, filtered 300-5000 Hz). Epidural EEG (filtered 0.5-300 Hz) and muscle tone EMG (filtered 10-100 Hz) were first passed through RA16LI preamplifier (TDT), then, together with microwire data, amplified by PZ2 amplifier (TDT) and synchronized with a video recording system RV2 (TDT). EEG and EMG data were recorded at a sampling rate of 256.99 Hz, and resampled offline at 1,000 Hz using custom written Matlab scripts (version 9.6.0 R2019a, The MathWorks, Natick, MA, USA). Narrow-band (0.1-0.2 Hz FWHM) notch filters were applied to the EEG signal to remove potential line noise at 50Hz and its harmonics, as well as at 7Hz (to remove noise at that frequency identified in some sessions). Spike sorting was performed offline with “wave_clus” (Quiroga et al. 2004), using a detection threshold of 5 SD above the median, and applying automatic superparamagnetic clustering of wavelet coefficients followed by manual refinement based on consistency of spike waveforms and inter-spike-interval distribution.

### Auditory stimulation

All experiments were conducted in a soundproof chamber attenuating 55dB of ambient noise (H.N.A, Kiryat Malachi, Israel). Sounds were synthesized in Matlab, transduced to voltage signal (195kHz sampling rate, RZ6, TDT), passed through a zero-amplification speaker amplifier (SA1, TDT), and played free field via a magnetic ultrasonic speaker (MF1, TDT) mounted 55 cm above the center of the cage (Figure 1B). We obtained an estimate of the sound intensity by placing a high-quality ultrasonic calibration microphone model 378C01 (PCB Piezotronics, Depew, NY, USA) at the center of the cage floor directly below the speaker (although the stimulus intensity experienced by the animal would vary depending on their precise position in the cage). During experimental sessions, rats were presented with 500ms-long 40Hz click-trains (20 clicks) at three different intensity levels (76, 82, 93 dB SPL) and a sham stimulus. Preliminary analysis showed no significant effects of stimulus intensity on neuronal responses, thus trials of the three intensities were pooled together in all subsequent analyses. The stimuli were separated by intervals of 1±0.4 seconds, and played in blocks of 120 repetitions (i.e. 360 stimuli), lasting 12 minutes each.

### Experimental design and anesthesia

Experiments started at least 7 days after surgery, once animals showed signs of full recovery. Two additional days allowed habituation to tethered recording and auditory stimulation (1 – 5 hours). On the day of the experiment, animals were placed in the recording chamber with wire bundles running to a counterbalanced anchor at the top, along with the IV catheter which was attached to a computer-controlled syringe pump (Chemyx Fusion Touch Series, Chemyx Inc., Stafford, TX, USA). Animals were gradually anesthetized with propofol, infused at a rate of 100 to 900-1200 mg/kg/min, to produce partial behavioral suppression (sedation) followed by complete behavioral unresponsiveness (Hudetz et al. 2011) (Figure 1C). The entire recording session lasted around 100 minutes (mean±SD: 105±26 minutes) depending on variability in response to anesthetic drug infusion and the number of blocks needed to achieve behavioral unresponsiveness.

At the end of each block after the animal was evaluated behaviorally (below), the infusion rate was increased. We estimated the instantaneous time-varying plasma propofol concentration (in mcg/ml) using a pharmacokinetic two-compartment model with Michaelis-Menten elimination (Ihmsen et al. 2002), based on the known rates of propofol infusion over time (Larsson and Wahlström 1994; Ihmsen et al. 2002; Banks et al. 2018).

### Behavioral assessment

Throughout the experiment and propofol infusion, intermittent behavioral assessment was performed every 12 minutes between blocks of auditory stimulation. This assessment was done to estimate the behavioral responsiveness as a function of the propofol levels of each animal/time (Church and Shucard 1987; Devor and Zalkind 2001; Banks et al. 2018; McKinstry-Wu et al. 2019; Du et al. 2020). Assessment included a battery of behavioral tests using standardized procedures (Devor and Zalkind 2001; Pillay et al. 2011): turning on back to assess for loss of righting reflex (LORR), tail pinch and hind-foot pinch to assess long spinal reflex, fore-paw pinch to assess short spinal reflex, dropping saline on eye for corneal reflex, touching whiskers for whisking reflex. Loss of consciousness (LOC) was operationally defined as loss of righting reflex (LORR) (Imas et al. 2005; Alkire et al. 2007; Laureys and Tononi 2011). Following LORR, some animals continued to exhibit responsiveness to some behavioral tests such as paw pinch before full loss of responsiveness (LOR) was achieved.

### EEG analysis

For each session, one of the EEG channels was selected for further analysis. EEG data were segmented into 1.5-second segments around each click-train stimulus, and grouped into the same 120-stimuli epochs used in the rest of this study. Noisy EEG segments (e.g. due to movement artefacts) were removed (5.4% of segments). Next, power spectra were calculated for each segment, averaged (mean) across all segments within a 120-stimulus block, and smoothed in the frequency domain by convolution with a Gaussian function (sigma = 1Hz). A peak in the spectrum in the alpha (7-15Hz) frequency band was identified by identifying peaks in that frequency range with prominence > 5% of peak power (red cross in Fig 1D & Supplementary Fig 3). The onset of alpha enhancement was defined as the first occurrence of at least two consecutive epochs showing such alpha peaks (red vertical line in Fig 1F & Supp Fig 1D).

### Histology

After completing data collection, rats were deeply anesthetized (5% isoflurane) and transcardially perfused with saline followed by 4% paraformaldehyde. Brains were extracted from skull, submerged in 4% paraformaldehyde for at least one week, and sectioned in 50-60 µM serial coronal sections using a vibrating microtome (Leica Biosystems, Israel). Electrode positions were determined by co-registering Dil traces in wet sections with subsequent fluorescence Nissl staining.

### Identification of responsive neuronal clusters and auditory response components

All analyses were carried out using custom-made Matlab procedures (2019a and Statistics Toolbox; MathWorks inc., Natick, MA, USA). For each neuronal cluster separately, we detected time intervals with significant increased spiking activity in responses to auditory stimuli (Sela et al. 2020), by comparing every millisecond in the [0-600ms] interval after stimulus onset (the 500ms of stimulus + 100ms following it) with all 500ms periods of pre-stimulus baseline activity, using a one-tailed Mann-Whitney test. We corrected for multiple comparisons using the False Discovery Rate (Benjamini and Yekutieli 2001) with base alpha of 0.001. Responses shorter than 5ms were excluded, and undetected intervals shorter than 5ms that proceeded and followed responses were categorized as misses and bridged with adjacent intervals. Each unit spike raster matrix was divided into epochs of 120 trials (∼4 minutes) and each epoch was evaluated separately. If one epoch had significant response the unit was considered responsive. We divided our responses into the following (non-mutually exclusive) classes:

Units with onset and offset responses. To identify onset responses, we focused on the time interval [9-25ms] after stimulus onset. Units with a significant response in at least one epoch in this time interval were labeled as ‘onset units’. Similarly, units that showed a significant response in a least one epochs in the interval [11-100ms] after stimulus offset were labeled ‘offset units’.

Units with 40Hz locking. We identified units that responded in a time-locked manner to 40Hz click-trains using inter-trial phase coherence (ITPC)(Tallon-Baudry et al. 1996):

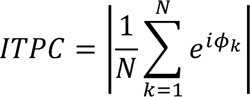

where N is the number of action potentials and ɸk is the phase of the action potential, k, within a recurring 25 ms (i.e. 40Hz) time window. Units with a 40Hz locking response were those that exhibited p < 10^-7^ (Zar 2010) and ITPC > 0.15.

### Analysis of ongoing and stimulus-driven dynamics during gradual anesthesia induction

For each unit and each 120-trial epoch separately, we quantified six components of the ongoing/auditory-evoked responses: (1) Baseline firing rate: for all the units that had a significant response to the auditory stimuli, we calculated the average firing rate across all the 500ms intervals prior to stimulus onset. (2) Onset response amplitude: for all ‘onset units’, we calculated the average firing rate within the [5-29]ms interval following stimulus onset, normalized (by subtraction) to the baseline firing rate of that bin. (3) Onset latency: for all ‘onset units’, we calculated the first significant response in the [1-100]ms interval following stimulus onset. (4) Post-onset silence: for all ‘onset units’, we calculated the degree of post-onset silence as follows. We computed the peri-stimulus time histogram (PSTH) in the [30-99ms] interval after stimulus onset, applied minimal smoothing to the PSTH via convolution with a Gaussian kernel (σ=2ms), and identified the duration of reduced firing as the interval between the first and last time points where firing was below 50% of that epoch’s baseline. (5) 40Hz locking: for each unit with 40Hz locking, we quantified the locking in that epoch by summing the phase values corresponding to each spike in the [150- 500] ms interval following stimulus onset. (Any firing that is not locked to the click-train does not contribute to this sum):

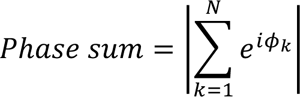

Where N is the number of action potentials and ɸk is the phase of the action potential, k, within a recurring 25 ms (i.e. 40Hz) time window. This measure was chosen over ITPC since it includes within it the effect of firing rate. A separate quantification using spike time tiling coefficient (STTC) (Cutts and Eglen 2014) yielded almost identical results. (6) Offset response: for all ‘offset units’ we calculated the magnitude of the offset response by taking the average firing rate in the [11-100ms] interval after stimulus offset, and normalizing (by subtraction) to the baseline firing rate of that epoch.

To go beyond variability in individual propofol concentrations and differences between individual unit firing, each of the six components above was normalized and expressed as percent relative to the maximum of that unit in that session (e.g. maximal baseline firing rate, maximal 40Hz locking). Along the same line, propofol concentration for each epoch was normalized and expressed as percent relative to the propofol concentration needed to achieve LORR in that session. Whilst the concept of “depth of anesthesia” is complicated, involving often subjective assessment of several behavioral measures or responsiveness, autonomic and brainstem functions, or approximations using various algorithms based on scalp EEG, we operationalize this LORR-normalized-propofol concentration as a quantitative measure of depth of anesthesia. From here we calculate the average gradient (change in neuronal activity as a function of change in propofol concentration) for each measure of interest (for example, -0.32 for decrease in baseline firing rates in Figure 3B corresponds to a change of -0.32% from original baseline firing with each 1% increase in normalized propofol concentration, or 32% decrease by the moment of LORR and 38% decrease by the end of recording at 120% normalized propofol concentration).

### Modeling neuronal dynamics with linear vs. step-wise descent functions

We estimated the fit of linear vs. step-wise functions to each component of the baseline/auditory-evoked response of each unit separately (if it had a least 10 epochs, i.e. including all sessions except Session #1 – see Table 1), using a leave-one-out method. We fitted both a linear function, a flexible timing step function with three free parameters: value before step, value after step, and timing of the step point that can be anywhere across the data, step function with a fixed inflection point at LORR and a step function with flexible timing but constrained to be the same for all neuronal clusters per animal. We used n-1 samples (epochs), and calculated the mean-square-error (MSE) of the model to the left-out sample. The error of each model was the mean over all possible leave-one-out combinations, normalized by dividing the MSE by the variance.

**Table 1.**
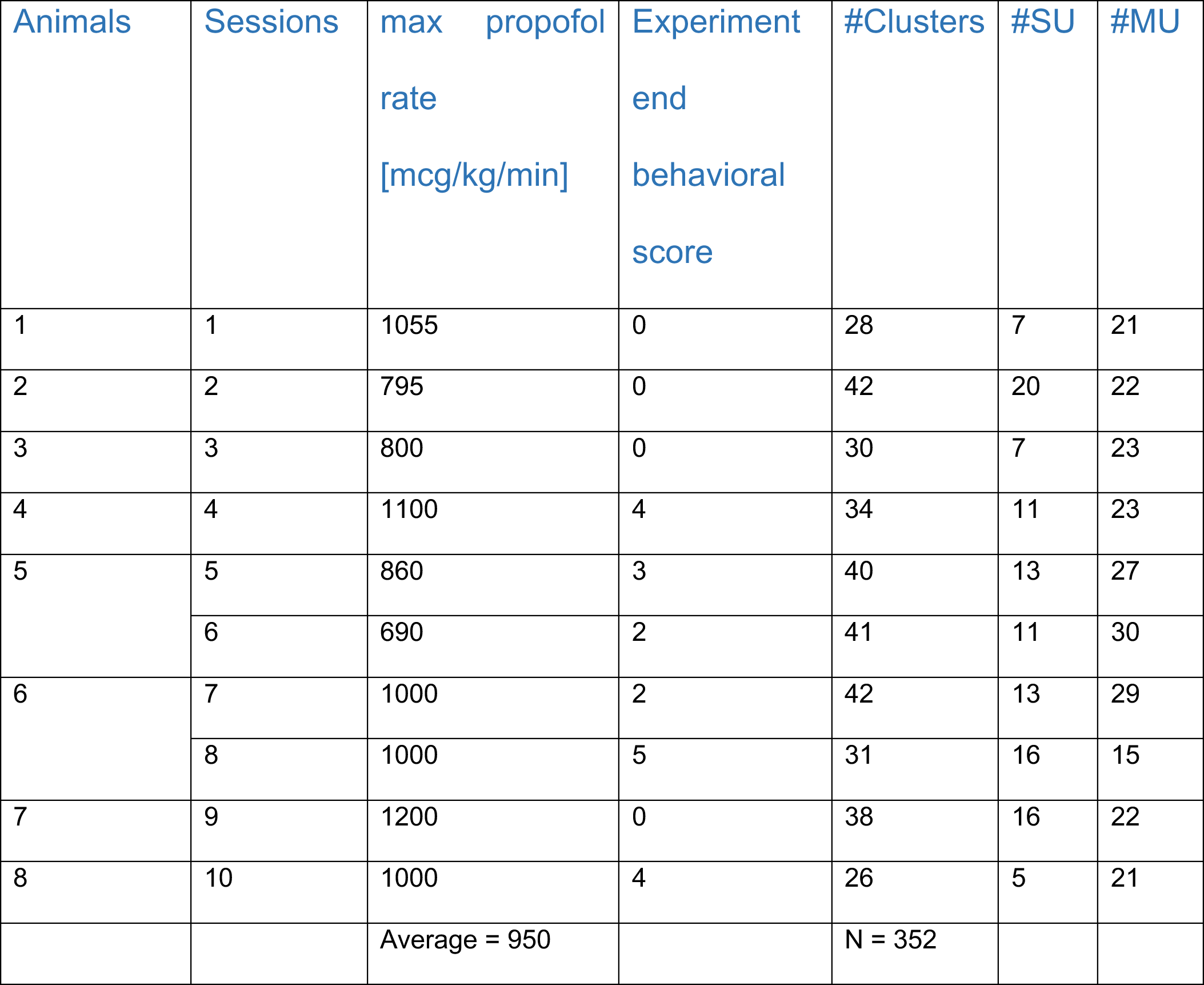
Data acquisition details. Experiments were conducted in ten animals, three of which participated in two separate experimental sessions. Two animals’ data were discarded due to technical problems with anesthesia or because histology revealed that electrodes did not reach PAC. # Clusters refers to all neuronal clusters identified in spike soring (n=352). Propofol infusion rates in each experimental session were increased until reaching stable LOR, with some variability across experiments (average maximum = 950 mcg/kg/min). Behaviorally, some rats were still showed some signals of responsiveness after LORR (mostly response to toe pinch).

### Statistics

To determine the statistical significance of each linear model fit to the real neuronal data, we employed a non-parametric approach where, for each analysis of interest, we compared the distribution of real data fits to fitting randomly shuffled data via Monte-Carlo Permutation Test. Shuffling re-sorted the order of the values for each unit between 120-trial epoch according to a random permutation such that the shuffled data was no longer consecutive in time and disrupted the relation between propofol levels and neuronal measures. We continued to compute the measure of interest (1-6 described in previous section) using the exact same procedure as for the real data. This procedure was repeated across 10,000 iterations to create the null distribution of each measure of interest. The P-value associated with each real measure was computed by examining the percent of random iterations that showed an average measure equal or greater than the single median observed in real data.

## Results

We tested whether PAC neuronal responses during gradual anesthesia are primarily determined by anesthesia concentration vs. stepwise change around loss of behavioral responsiveness. Anesthesia was slowly induced by propofol infusion through intravenous jugular catheters (from 100 to 900-1200 mcg/kg/min), combined with intermittent behavioral assessments. PAC spiking activity was recorded in response to 500ms-long auditory 40Hz click-trains. We conducted ten experimental sessions, lasting 60 minutes each, in eight freely behaving rats (Figure 1). Every 12 minutes, auditory stimulation was paused and the animal was assessed behaviorally for righting and other (e.g. corneal, paw withdrawal, whisking) reflexes, after which the rate of propofol infusion was increased. Experiments terminated upon complete cessation of all reflexes, at least 12 minutes after LORR.

In addition to behavioral assessment of LORR, we identified the appearance of enhanced alpha (7-15Hz) activity in the epidural EEG, representing an electrophysiological correlate of LOC (Purdon et al. 2013; Krom et al. 2020) (Figure 1D & Supplementary Figure 3). In all cases, EEG alpha activity appeared at significantly (p< 0.008 via Wilcoxon signed rank test) lower propofol concentrations than LORR (Figure 1E), around mean (±SD) normalized propofol concentration of 20.5% (±27.2%). In all cases the peak frequency of such EEG activity drifted downwards as propofol concentration increased (median reduction of 0.31 Hz per epoch).

Neuronal activity was recorded continuously with arrays of microwire electrodes targeting the right PAC. We isolated 352 neuronal clusters (119 single units, 233 multiunit clusters; Table 1), of which 243 units (69%) had a significant response to the 40Hz click-train auditory stimuli. To verify the focus on PAC activity, we selected a subset of 195 auditory-responsive clusters that that were histologically localized to PAC (Figure 2A,B) and exhibited a significant onset response with latency of 9-25ms (Figure 2C, Methods) for subsequent analysis. We refer to these data throughout as PAC despite the lack of systematic tonotopic mapping (see also Discussion).

**Figure 2.**
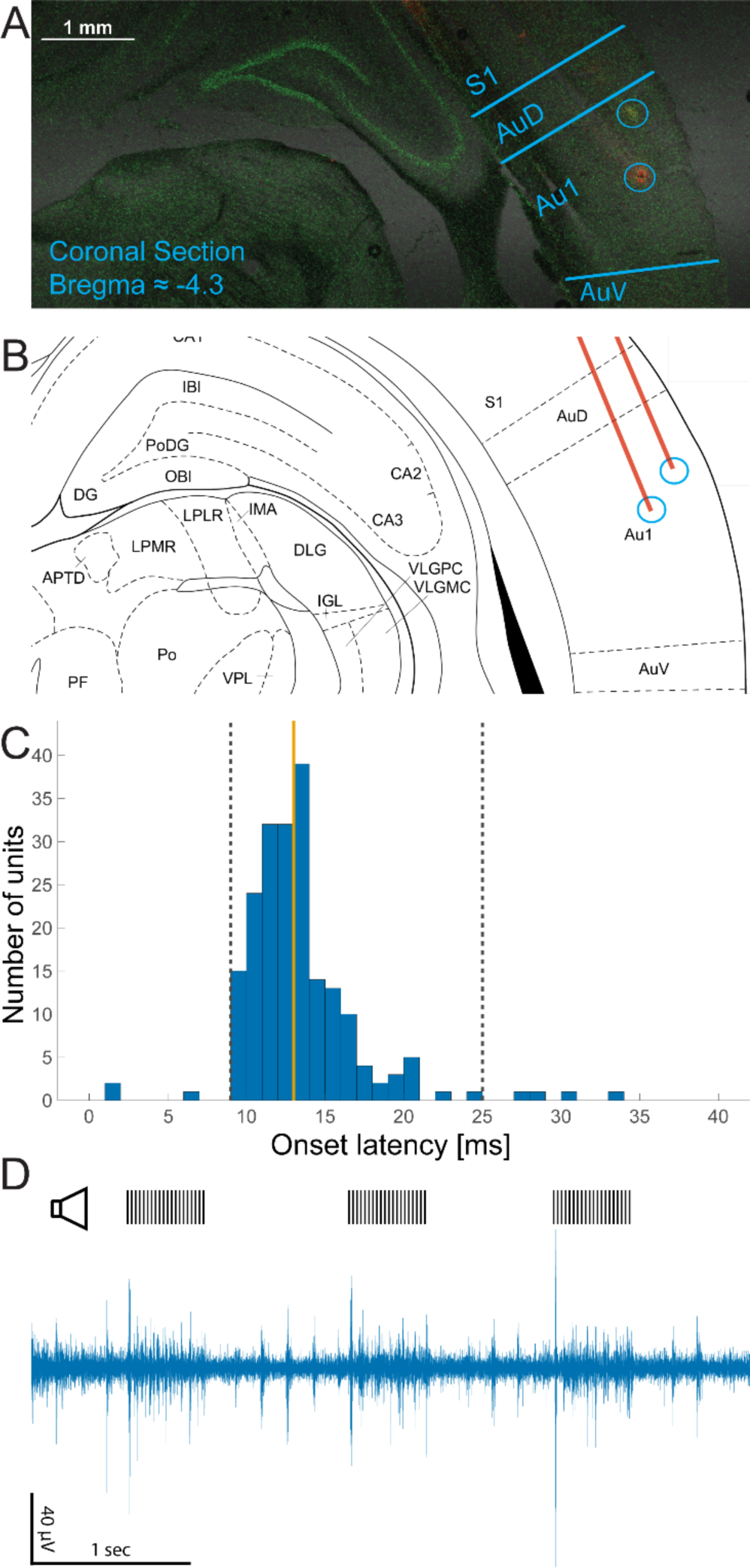
Auditory cortex targeting. (A) Representative coronal brain section (∼4.3mm posterior to Bregma) showing histological verification of electrode positions in auditory cortex (circles show distal microwire tips with DiI). (B) Corresponding coronal section from Paxinos atlas (Zhang et al. 2015). (C) Distribution of onset latencies across all units with a significant onset response (n=221). Orange line marks median latency (13ms), dashed black lines mark units with latencies within [9- 25ms] (88%) used for subsequent analyses. (D) Recorded signal from channel (bandpass 300- 5000Hz), upper black line represents the auditory stimuli (40Hz click train).

### PAC Neuronal units exhibit different response components to sound stimuli, which deteriorate going from wakefulness to deep anesthesia

Figure 3A provides a representative example of neuronal activity during the experiment. The neuronal responses consisted of multiple components including onset responses (<30ms), post-onset silence (30-100ms), locking to the 40Hz click-train stimulus (150-500ms), and occasional offset responses (during the 100ms after stimulus termination). Accordingly, neuronal clusters could be divided to partially-overlapping subgroups exhibiting onset responses (n=195), post-onset silence (n=188), locking to the 40Hz click-train stimulus (n=118), and offset responses (n=73).

**Figure 3.**
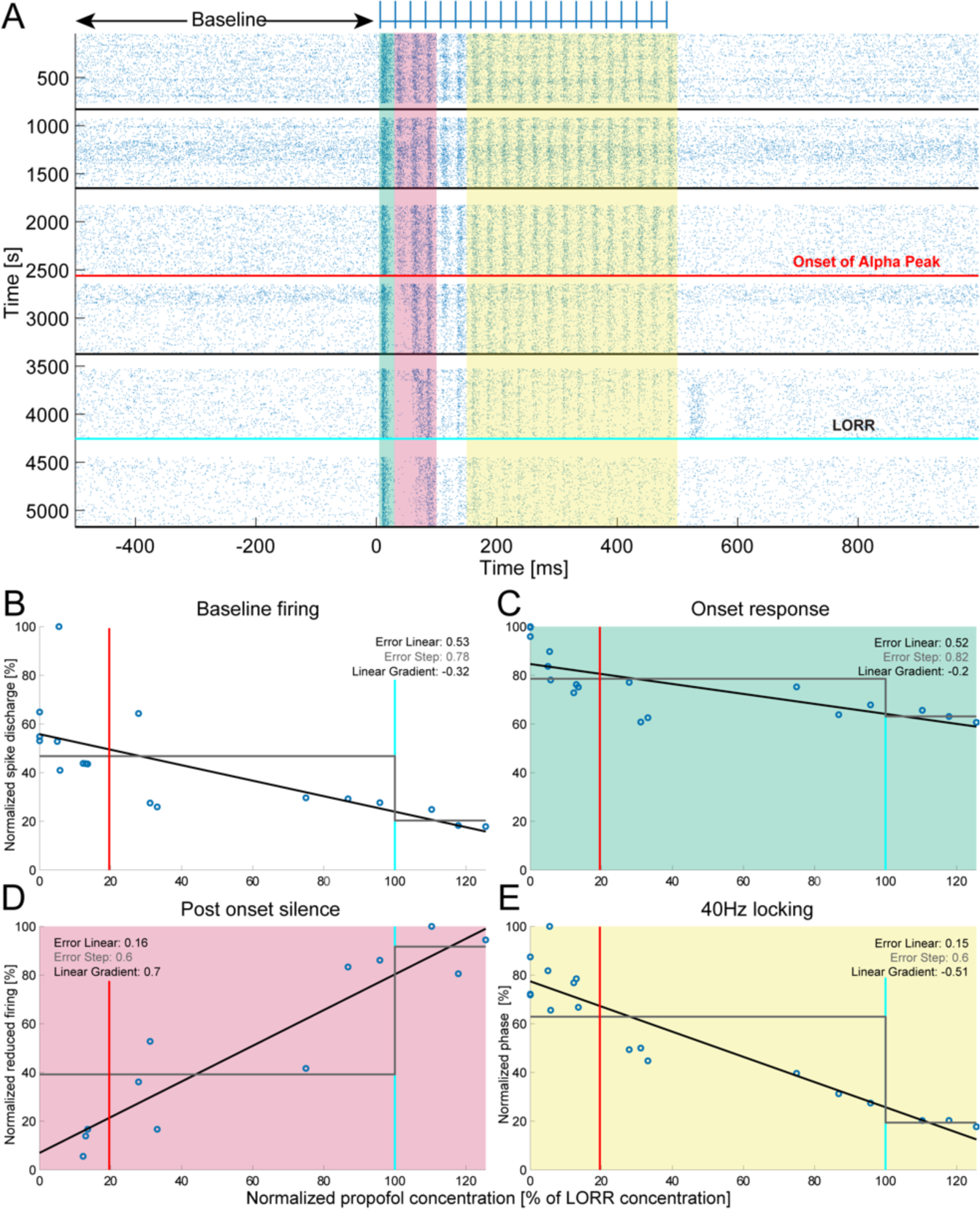
Representative neuronal responses to 40Hz click-trains in rat PAC during gradual deepening of propofol anesthesia. (A) Spike raster plot during the entire experiment (rows represent trials in seconds 1-5172, propofol infusion is increased from top to bottom). Blue ticks on top mark individual clicks in the click-train stimulus. Horizontal black lines mark intermittent behavior assessments performed every 12 minutes. Bottom-most black line shows termination of experiment upon complete loss of responsiveness including reflexes, while the cyan line above it marks LORR. Horizontal red line shows onset of alpha enhancement. Green shading, time interval for response onset [5-29]ms. Red shade, time interval for post-onset silence [30-99ms]. Yellow shade, time interval for ‘late locking’ to clicktrain [150-500]ms. White rows of discontinuity in raster plot represent the times of behavioral assessment when no sounds were delivered. For each component of interest (baseline firing, onset response, post-onset latency, and 40Hz locking) we calculate the response properties, (normalized to the maximum response) as a function of the propofol concentration (expressed as percent of dose eliciting LORR). (B-E) Dynamics of spike discharge rates, for the same neuronal cluster shown in A, in distinct temporal intervals around the auditory trials, separately for (B) Baseline firing [-500 0]ms, (C) Onset response, green highlight, (D) Post-onset silence, red highlight (E) Late locking to 40Hz click-trains, yellow highlight. Blue circles in each panel show the average measure of interest in 4-minute epochs (corresponding to 120 trials). Black lines show linear fit whereas grey lines show step-wise fits with inflection point in LORR (Methods). Vertical red and cyan lines mark onset of alpha enhancement, and LORR, respectively. Insets on upper-right depict error quantifying each model’s fit and linear gradient. Note that all measures are modeled better by gradual linear

Neuronal activity robustly deteriorated during the progression of the experiment. When comparing the first and last 12-min periods (corresponding to drug-free wakefulness vs. deep anesthesia without behavioral responsiveness, respectively) we found that mean baseline firing rates decreased by 57±1.5 % (Mean ± S.E.M, n=243 clusters, p<10^-41^, Wilcoxon signed-rank test), mean onset response magnitude decreased by 31.5±2.2% (n=195 clusters; p<10^-24^, Wilcoxon signed-rank test), mean onset latency modestly increased by 7±1.18% (n=195 clusters; p<10^-11^ via Wilcoxon signed-rank test), the mean post-onset silence increased by 30.1±2.24% (n=195 clusters; p<10^-23^ via Wilcoxon signed-rank test), and mean late locking to 40Hz click-trains decreased by 61±2% (n=118 clusters; p<10^-21^ via Wilcoxon signed-rank test). Offset responses, whenever present, increased in magnitude upon anesthesia by 30±9.8 % (n=73 clusters; p=0.02 via Wilcoxon signed-rank test) but their occurrence was sporadic.

### Changes in neuronal activity are gradual rather than abrupt

To test whether neuronal activity changed in closer association with gradual anesthesia depth vs. abruptly around the point of loss of responsiveness, we quantified each component-of-interest in 4-min epochs (120 trials each), and compared the goodness-of-fit of linear vs. stepwise functions in modeling the dynamics during the descent to anesthesia (Figure 3B-E). In the representative cluster shown in Figure 3, gradual linear decay constituted a better fit (higher explained variance) for the dynamics of most activity components (baseline firing, onset response, post-onset silence, and 40Hz locking). A similar pattern of gradual decay was observed in the activity and response dynamics of many other clusters and animals (Supplementary Figure 1).

We proceeded to compare the goodness-of-fit of linear vs. step-wise models across the entire dataset by calculating, for each neuronal cluster and response component separately, the difference between the errors of linear vs. step-wise models (Table 2 and Figure 4). For most components of interest (baseline firing, onset response magnitude, post-onset silence, and 40Hz locking) the distribution of differences was skewed towards negative values, indicating better fit of linear (gradual) models. We assessed the statistical significance of these differences by comparing, via Wilcoxon signed rank tests, the distributions of error values between linear and step-wise functions. We found that in all the four components-of-interest, the difference was highly significant for all models (Table 2), indicating that linear (gradual) models better fit changes in neuronal activity in our data than step-wise models.

**Figure 4.**
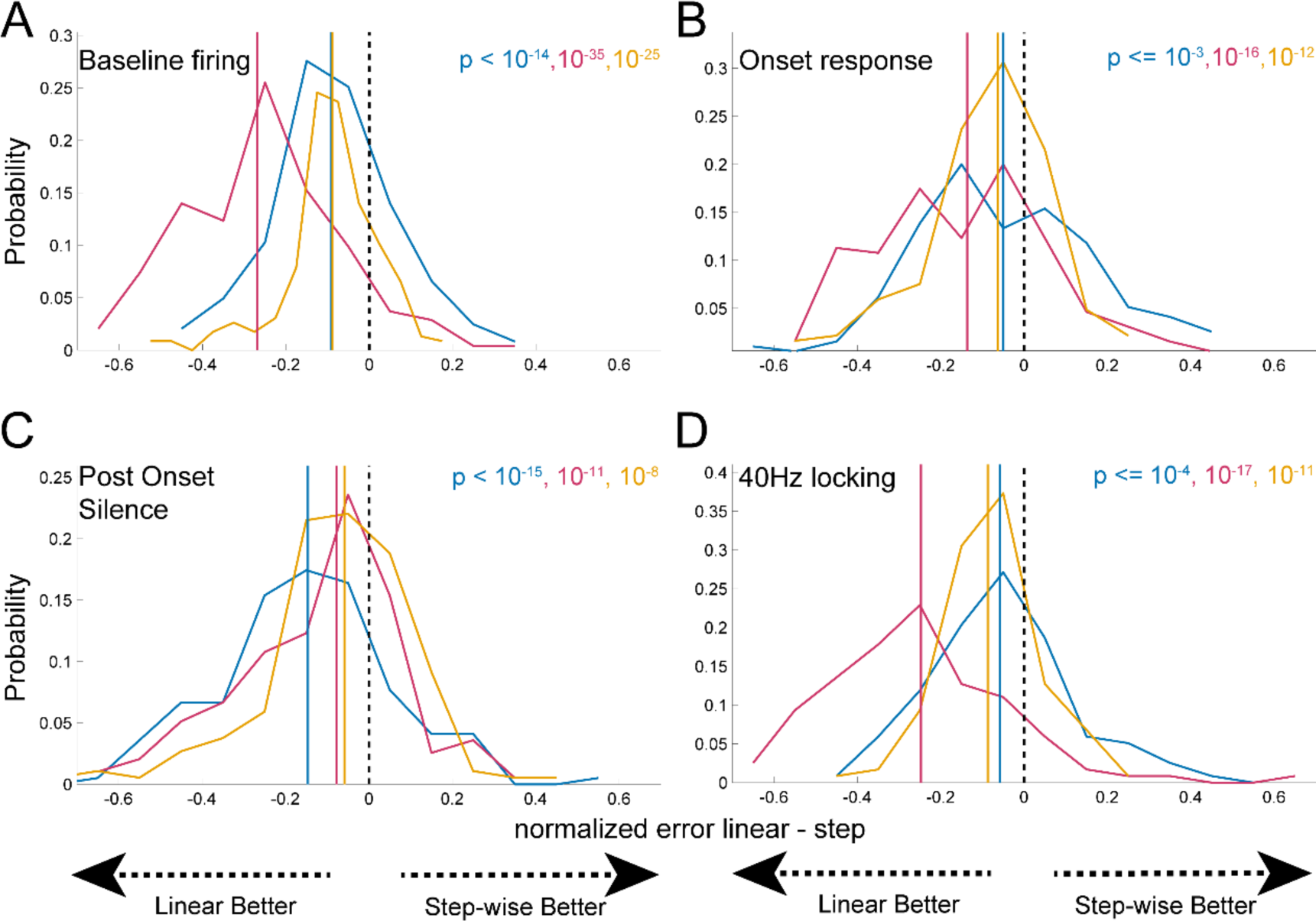
Linear models better explain changes in ongoing and stimulus-driven neuronal activities than stepwise models upon deepening of anesthesia. Each panel represent the subtraction between errors (goodness of fit) of linear vs. step-wise models across the entire neuronal population. (A) Baseline firing (N=243). (B) Onset responses (N=195). (C) Post-onset silence (N=188). (D) 40Hz locking (N=118). Black dashed lines, zero difference (linear and step-wise models equally fit the data), each curve represent the distribution of the subtraction between the linear and a step model, and each corresponding line is the median of this distribution. Blue curve and line: step model with flexible timing; orange curve and line: step model with flexible timing but fixed for all clusters per animal; red curve and line: step model with fixed timing at LORR. Note that all distributions are skewed to the left, showing that linear fits match the data better than any step wise fits for all response elements. P-values in upper-right corner of each panel mark the statistical significance of linear models fitting the data better than step-wise models (via Wilcoxon signed rank tests, Methods).

**Table 2.** Comparison of linear vs step-wise models to fit the data. Negative values imply linear fit is best. N is the number of units included. P-values by Wilcoxon signed rank test on errors of linear vs relevant stepwise model.

|  |  | Free<br>inflection<br>point |  | Inflection at<br>LORR |  | Fixed<br>inflection per<br>animal |  |
| --- | --- | --- | --- | --- | --- | --- | --- |
|  | N | median | p | median | p | Median | p |
| Baseline<br>firing | 243 | -0.09 | $<10^{-14}$ | -0.27 | $<10^{-35}$ | -0.09 | $<10^{-25}$ |
| Onset<br>response<br>magnitude | 195 | -0.05 | $<10^{-3}$ | -0.14 | $<10^{-16}$ | -0.06 | $<10^{-12}$ |
| Post-onset<br>silence | 188 | -0.15 | $<10^{-15}$ | -0.08 | $<10^{-11}$ | -0.06 | $<10^{-8}$ |
| 40Hz<br>locking | 118 | -0.06 | $<10^{-4}$ | -0.25 | $<10^{-17}$ | -0.09 | $<10^{-11}$ |

### Sensitivity to propofol concentration varies between response components

To better characterize the association between increases in propofol and degradation of neuronal responses, we examined the slopes (gradients) of the linear models for each response component separately (Figure 5). The median slope (gradient) values were -0.41 for baseline firing, -0.24 for onset response magnitudes, +0.3 for post-onset silence, and -0.41 for 40Hz locking, respectively (figure 5). These values represent a robust change of around 20-40% in each response component during the entire descent from drug-free wakefulness to LORR, and represent reduction of 35-60% when continuing to deeper depths of anesthesia (150%). In contrast, when progressing from wakefulness to the onset of alpha enhancement (at an average of 20.5% normalized propofol concentration), possibly correlated with undisturbed loss of consciousness(Purdon et al. 2013), the response components change approximately 5- 8%. We assessed the statistical significance of these slopes (compared to no linear deterioration) by comparing the real data to surrogate data where the order of bins was randomly shuffled in time to eliminate the link to propofol concentration using a Monte-Carlo Permutation Test (Methods). We found that the linear gradients fitting the real data were significantly smaller (or larger in the case of post-onset silence that increased with anesthesia) than shuffled data (all p<0.0001).

**Figure 5.**
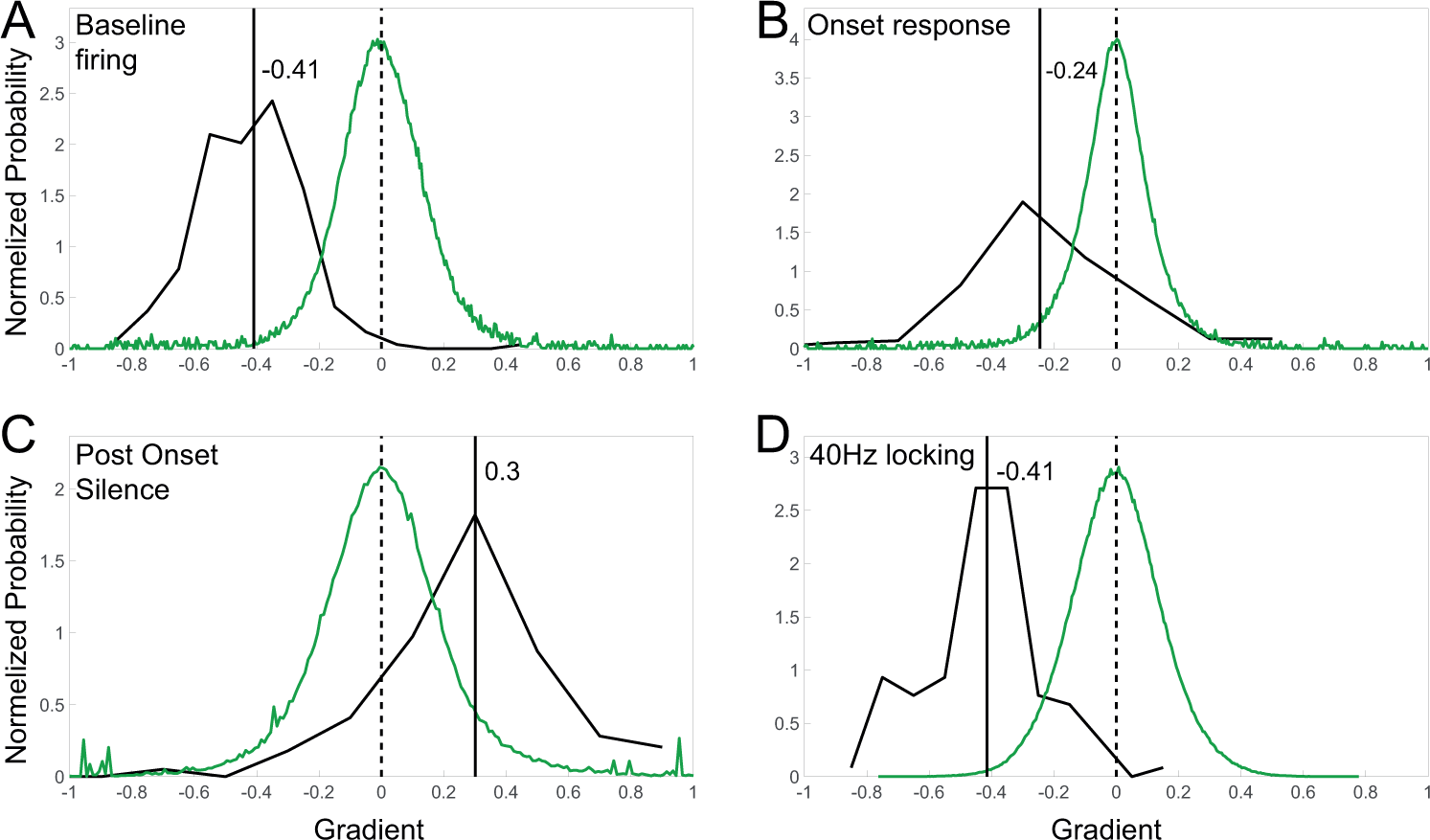
Rates of linear changes in neuronal activity as a function of propofol concentrations upon deepening of anesthesia. Each panel represents the rate (gradient) of linear changes for each measure of neuronal activity as a function of normalized propofol concentration, across the entire neuronal population. (A) Baseline firing (N=243). (B) Onset responses (N=195). (C) Post-onset silence (N=188). (D) 40Hz locking (N=118). Black curves, distribution of gradients. Black solid vertical lines, median gradient. Green curves, distribution of 10,000 surrogate gradients obtained from shuffled data. Black dashed lines, zero.

In addition to the main components-of-interest, the magnitude of offset responses, as well as the latency of onset responses, increased in association with propofol concentration. The increase in onset latency with propofol concentration was significant but modest (median gradient = 0.06, representing a 6% increase in latency from drug-free wakefulness to propofol concentration that achieve LORR, p=<0.001 via Monte Carlo Permutation Test, or 9% if continued to deeper depths of anesthesia). In this case a step-wise function better fit the data than a linear model (p < 0.01). Offset responses, whenever present (n=73), grew stronger during deepening of anesthesia, although there was large variability between neuronal clusters and epochs. This increase was statistically significant (median gradient = +0.14, representing a 14% increase in offset response magnitude from drug-free wakefulness to propofol concentration that achieve LORR, p=<0.001 via Monte Carlo Permutation Test). Similarly to the nearly all other response components, offset responses were better modeled by linear rather than by stepwise functions (p = 0.0019, via Wilcoxon signed rank test). We note however that the result was less consistent across units, and that the number of the units with offset response was smaller (n=73).

To understand which aspect was more sensitive to propofol we examined the sub population of units for which all the aspects of responses could be calculated (units with 40Hz locking, n=118). We found that the order from most to least sensitive was baseline firing (median: -0.41) ≈ 40Hz locking (median: -0.41) > post-onset silence (median: 0.32) > onset response (median: -0.2) > latency of onset responses (median: 0.06), and that these differences were significant (p = 10^-51^, Kruskal-Wallis test). This result confirms the order of sensitivity to propofol (figure 5) when including the full set of units. It is also worth noting that that gradients for all response components are mostly in the range of -0.2 to -0.4, representing a highly robust change of around 20- 40% in each response component during the entire descent from drug-free wakefulness to LORR.

## Discussion

### Neuronal auditory responses degrade gradually and linearly with increasing propofol concentration

Using gradual propofol anesthesia induction in rats while examining neuronal auditory responses to 40Hz click-train stimuli, we find that depth of anesthesia (propofol concentration) linearly degrades PAC activity. The main components of PAC activity including spontaneous firing rates, onset response magnitudes, onset response latencies, post-onset neuronal silence duration, and late-locking to 40Hz click-trains, gradually deteriorated in a dose-dependent manner with increasing anesthesia levels. Formal comparisons established that linear models constituted a better fit for PAC activity degradation than step-wise models (with either a-priori or post-hoc flexible timing for the stepwise decline, Figure 4). Moreover, such a flexible stepwise analysis did not prefer the time point of EEG alpha enhancement (Figure 1D-E & Supplementary Figure 3, often considered as an electrophysiological correlate of human LOC (Ching et al. 2010; Purdon et al. 2013)) over a gradual linear fit to the data. Thus, our gradual induction protocol successfully teases apart the effects of anesthesia depth vs. behavioral state changes (often confounded in many previous studies) and establishes that anesthesia depth (rather than abrupt changes around LORR or other time-points) is the key factor driving response changes in PAC.

Analysis of linear slopes (gradients) shows that during the full descent from drug-free wakefulness to deep anesthesia with behavioral unresponsiveness, PAC activity gradually deteriorates (Figure 5). Some response components (e.g. changes in baseline firing rates and late-locking to the 40Hz stimulus) were more sensitive than others to anesthesia depth and underwent larger deterioration with increasing anesthesia compared to other components (onset response magnitude and latency) that only exhibited modest changes (41% vs 6-24%). The relative invariance of onset responses to anesthesia is in line with other studies showing that responses to single brief events (or few events presented at a low frequency) such as auditory clicks, visual flashes (Wang 2018), or whisker deflections (Sharon and Nir 2018) are more preserved across behavioral states.

### Our results explain how studies comparing wakefulness and anesthesia can either show large degradation or modest changes

How do our results relate to previous studies comparing responses across wakefulness and anesthesia? A survey of the literature reveals a wide spectrum of reported results. Many studies across primary sensory cortices, both auditory (Castro-Alamancos 2004) and other modalities (Schwender et al. 1993; Capsius and Leppelsack 1996; Plourde 1996; Gaese and Ostwald 2001; Wang et al. 2018; Du et al. 2020) reported robust reduction (50-100%) in responses under anesthesia. Other studies from both auditory (Ishizawa et al. 2016; Banks et al. 2018; Nivinsky Margalit et al. 2020) and non-auditory (Heinke et al. 2004; Davis et al. 2007; Raz et al. 2014; Krom et al. 2020) primary sensory cortices showed more modest reductions (<21%) or even no reductions. We believe that the precise anesthesia drug concentration in each experimental condition is a key factor that could readily explain these discrepancies. Most previous studies contrasted two specific states (e.g. wake vs. deep anesthesia) and this is the case in all the studies listed above that showed large reductions in primary sensory cortical activity. In contrast, a number of the studies that showed only modest reductions in neuronal activity (Ferezou et al. 2006; Sellers et al. 2013), utilized sub-anesthetic levels of sedation. Our results recapitulate, within the same experiment, the spectrum of results reported in previous studies and resolve the apparent discrepancy in the literature by highlighting depth of anesthesia as a key factor dictating the extent of PAC degradation. Another important and often overlooked consideration is whether the ‘anesthesia’ condition is based on behavioral criteria such as LORR (failure to respond to a relatively aggressive external stimulus) or based on electrophysiological criteria such as EEG alpha enhancement (a measure correlated with LOC, occurring earlier). These two criteria are different in terms of their associated depth of anesthesia; surely, some conditions (e.g. natural sleep or deep sedation) are associated with LOC but would not exhibit LORR. Accordingly, our results show that PAC activity can exhibit degradation of either 20-40% or 5-8%, depending on whether the anesthesia condition is defined by LORR or LOC/alpha, respectively.

Our results join previous studies (Davis et al. 2007; Krom et al. 2020) demonstrating the effectiveness of the specific 40Hz click-train stimulus in comparing auditory responses across different states of anesthesia, and more generally as an index of auditory perceptual awareness. Accordingly, in human non-invasive electroencephalogram (EEG) studies, the locking to 40Hz click-trains often attenuates around anesthesia-induced LOC (Yli-hankala et al. 1994; Plourde et al. 2008; Purdon et al. 2013; Banks et al. 2018; Krom et al. 2020). Robust and relatively-abrupt changes in EEG 40Hz locking in these studies likely represent attenuated activity in association cortices beyond PAC, given that PAC responses only modestly contribute to total cerebral activity, and given the medial location of PAC that is less accessible to noninvasive scalp EEG. Indeed, a direct comparison between PAC and association regions upon anesthetic-LOC in human patients reveals modest PAC changes co-occurring with robust and relatively-abrupt changes beyond PAC (Galambos et al. 1981; Plourde et al. 2008; Purdon et al. 2013). Gradual and modest changes in primary sensory regions around LOC may lead to robust and abrupt changes downstream via non-linear threshold-like ‘ignition’(Krom et al. 2020). In states of unconsciousness and sensory disconnection, large neuronal populations seem unable to reliably synchronize with high-frequency stimulation (Leopold and Logothetis 1996; Moutard et al. 2015).

### Study limitations

A few study limitations should be explicitly acknowledged. First, we did not delineate auditory fields based on tonotopy. However, both histology and analysis of onset response latencies strongly suggest that the data represent PAC. Second, this study was conducted only with propofol (an archetypal GABA-agonist, (Sharon and Nir 2018)). While other anesthetics such as isoflurane and dexmedetomidine are also not associated with abrupt changes in PAC activity around LOC (Franks 2008; Brown et al. 2011), future studies should compare additional classes of anesthetic drugs such as ketamine and opioids. Third, behavioral testing was performed intermittently every 12 minutes. Future studies could track behavioral responsiveness more continuously with rotating cylinders or treadmills (Banks et al. 2018). Although differences in the precise drug or species may contribute to discrepancies in previous results, the fact that we recapitulate these differences within one experiment using a single species and drug argues that it may likely stem from depth of anesthesia.

To conclude, by employing a gradual induction of propofol anesthesia while examining auditory responses in the rat PAC, we were able to separate the effects of anesthesia levels from those of behavioral unresponsiveness. Our results show that propofol concentration constitutes the key factor in modulating responses in primary sensory cortex upon anesthetic-LOC, a factor that can explain seemingly conflicting results in the literature.

## Acknowledgements

We thank Maya Levin Arama for training in IV catheter insertion procedures, Becca Krom for illustration work, Israel Nelken for input, Hagai Bergman for comments on an earlier draft, and all members of the Nir Lab for discussions.

