## Supplemental Material for "Propofol Anesthesia Concentration Rather Than Abrupt Behavioral Unresponsiveness Linearly Degrades Responses in the Rat Primary Auditory Cortex"

### Supplementary Material

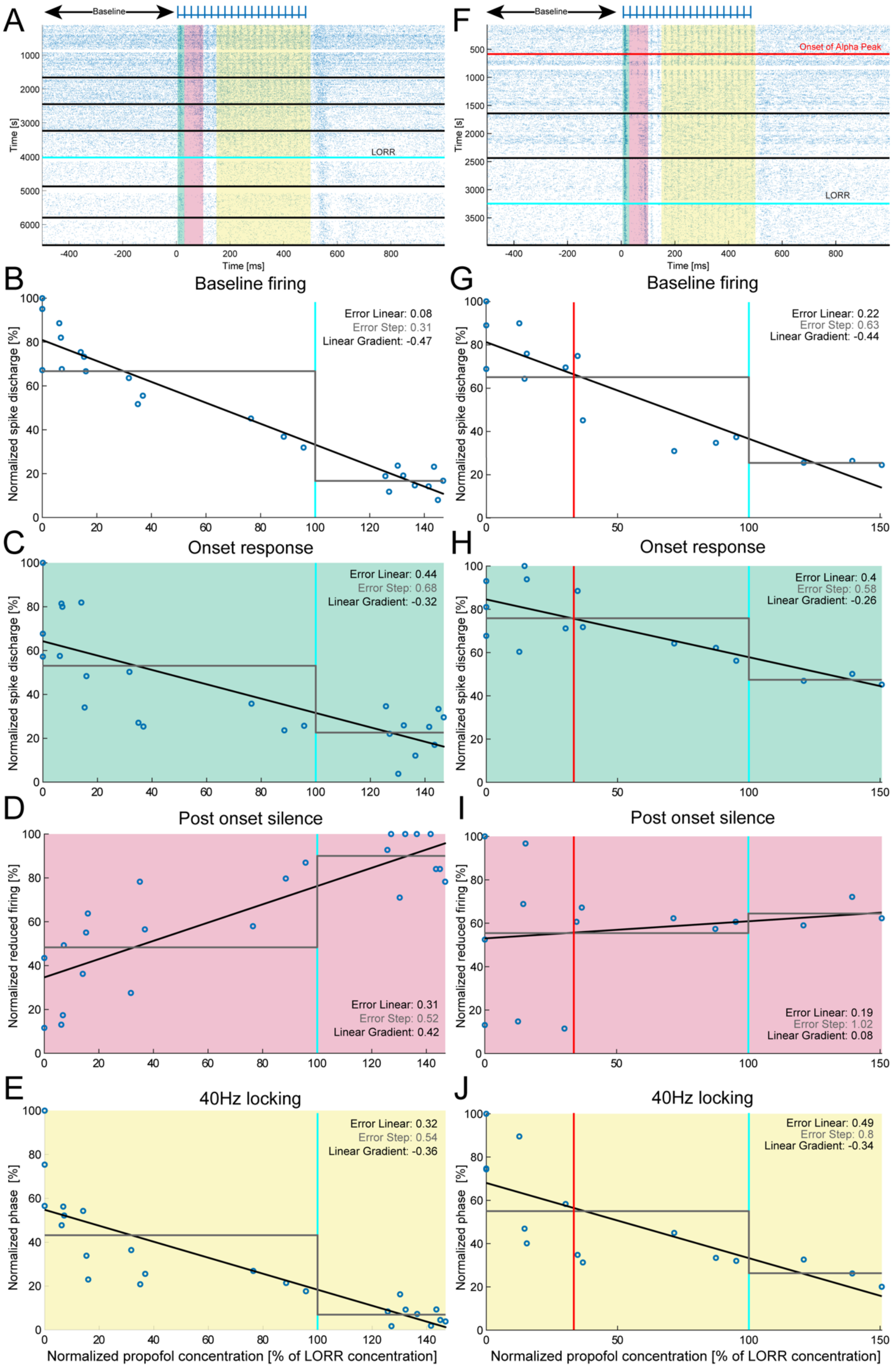

**Supplementary Figure 1. Additional representative PAC neuronal responses to 40Hz click-trains from 2 additional different rats during gradual deepening of propofol anesthesia.** (A,F) Spike raster plot during the entire experiment (rows represent trials in seconds 1-~6500, propofol infusion is increased from top to bottom). Blue ticks on top mark individual clicks in the click-train stimulus. Horizontal black lines mark intermittent behavior assessments performed every 12 minutes. Bottom-most black line shows termination of experiment upon complete loss of responsiveness including reflexes, while the cyan line above it marks LORR. Horizontal red line in (F) shows onset of alpha enhancement. Green shading, time interval for response onset [5-29]ms. Red shade, time interval for post-onset silence [30-99ms]. Yellow shade, time interval for 'late locking' to click-train [150-500]ms. White rows of discontinuity in raster plot represent the times of behavioral assessment when no sounds were delivered. For each component of interest (baseline firing, onset response, post-onset latency, and 40Hz locking) we calculate the response properties, (normalized to the maximum response) as a function of the propofol concentration (expressed as percent of dose eliciting LORR). (B-J) Dynamics of spike discharge rates, for the same neuronal cluster shown in A, in distinct temporal intervals around the auditory trials, separately for (B,G) Baseline firing [-500 0]ms, (C,H) Onset response, green highlight, (D,I) Post-onset silence, red highlight (E,J) Late locking to 40Hz click-trains, yellow highlight. Blue circles in each panel show the average measure of interest in 4-minute epochs (corresponding to 120 trials). LORR occurs at 100% Normalized propofol concentration by definition. Red vertical line in (H-J) marks appearance of alpha enhancement. Panels (A-E) refer to the single session (Session 10) where alpha enhancement could not be identified and so there is no red horizontal or vertical lines. Black lines show linear fit whereas grey lines show step-wise fits with inflection point in LORR (Methods). Insets on upper-right depict error quantifying each model's fit and linear gradient. Note that all measures are modeled better by gradual linear trends than by step-wise changes.

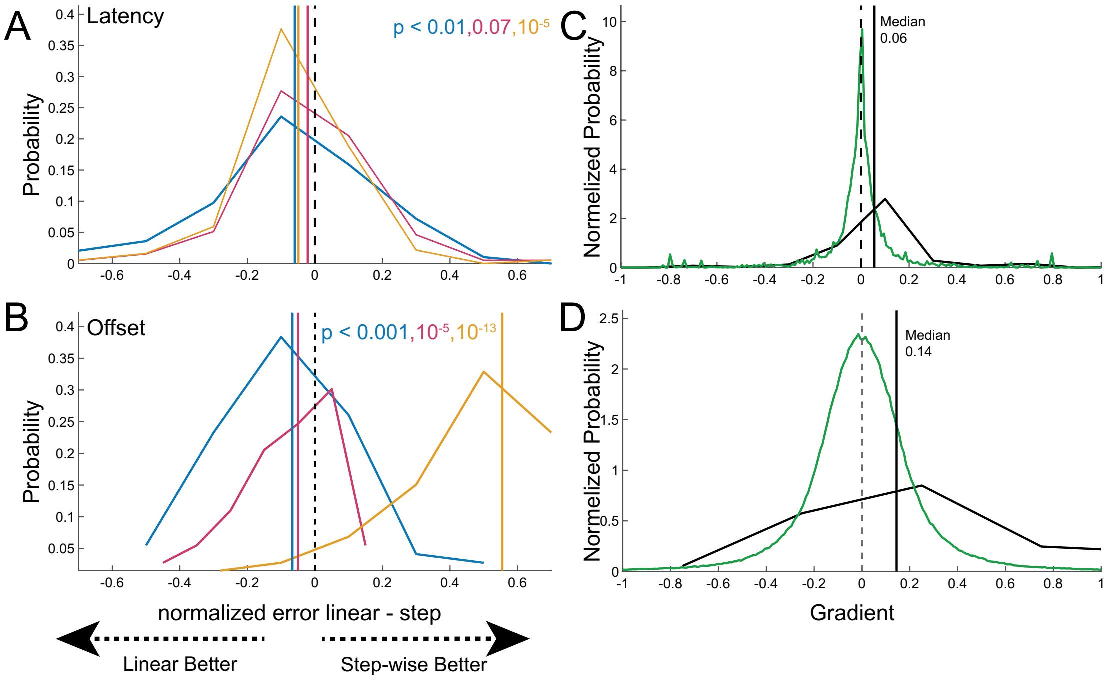


**Supplementary Figure 2. Latency and offset response, fitting decrease model and linear change.** Left panels represent the subtraction between errors (goodness of fit) of linear vs. step-wise models across the entire neuronal population**.** (A) Latency (N=195). (B) Offset responses (N=73). Black dashed lines, zero difference (linear and step-wise models equally fit the data), each curve represent the distribution of the subtraction between the linear and a step model, and each corresponding line is the median of this distribution. Blue curve and line: step model with flexible timing; orange curve and line: step model with flexible timing but fixed for all clusters per animal; red curve and line: step model with fixed timing at LORR. P-values in upper-right corner of each panel mark the statistical significance models fitting the data (via Wilcoxon signed rank tests, Methods). Right panels represents the rate (gradient) of linear changes for each measure of neuronal activity as a function of normalized propofol concentration, across the entire neuronal population. (C) Latency (N=195). (D) Offset responses (N=73). Black curves, distribution of gradients. Black solid vertical lines, median gradient. Green curves, distribution of 10,000 surrogate gradients obtained from shuffled data. Black dashed lines, zero.

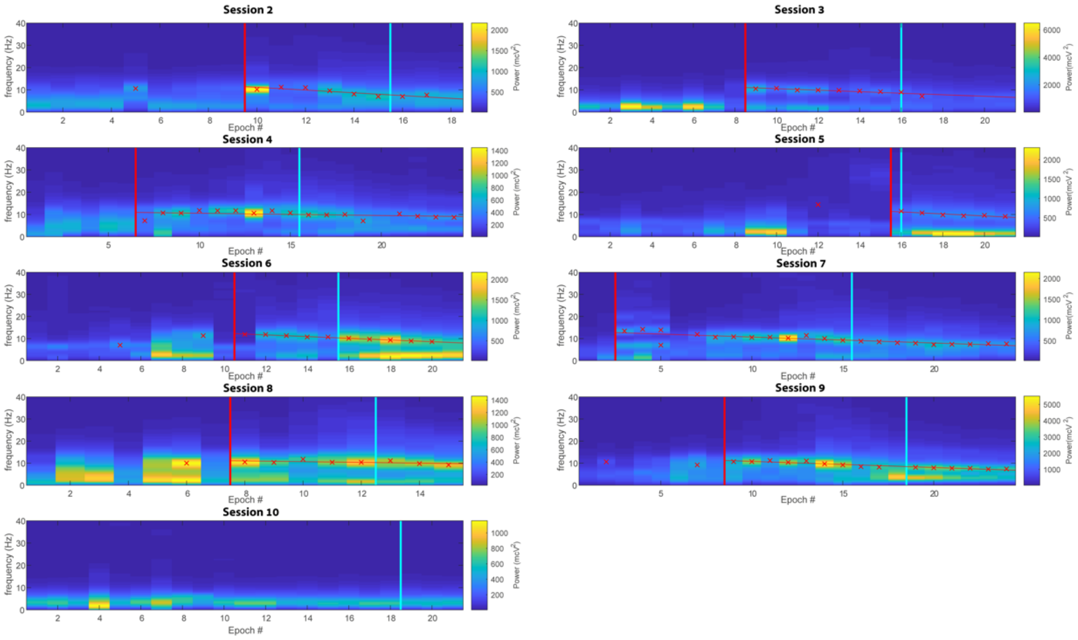


**Supplementary Figure 3. EEG time-frequency dynamics (spectrogram) per experimental session**. Each **panel** shows one experimental session (9/10 sessions where at least 10 epochs were available for EEG analysis). X-axis shows the temporal progression (120-trial epoch number). Y-axis shows frequency. Cyan vertical line shows moment of LORR. Red crosses show detected EEG alpha peaks (prominence >5% total power). Vertical red line indicates onset of alpha enhancement (Methods). Red line connecting asterisks shows the best linear fit through the alpha peaks after the onset of EEG alpha. Note that this line represents a consistent negative gradient (downwards drift in alpha frequency) with increasing propofol.
